# Transcriptional profiling of peripheral blood mononuclear cells identifies inflammatory phenotypes in ataxia telangiectasia

**DOI:** 10.1101/2023.10.05.561081

**Authors:** Nigel S. Michki, Benjamin D. Singer, Javier V. Perez, Aaron J. Thomas, Valerie Natale, Kathryn A. Helmin, Jennifer Wright, Leon Cheng, Lisa R. Young, Howard M. Lederman, Sharon A. McGrath-Morrow

## Abstract

Ataxia telangiectasia (A-T) is an autosomal recessive neurodegenerative disease with widespread systemic manifestations and marked variability in clinical phenotypes. In this study, we sought to determine whether transcriptomic profiling of peripheral blood mononuclear cells defines subsets of individuals with A-T beyond mild and classic phenotypes, enabling identification of novel features for disease classification and treatment response to therapy.

**Methods:** Participants with stable A-T (n=90) were recruited and compared with unaffected controls (n=15). PBMCs were isolated and bulk RNAseq was performed. Plasma was also isolated in a subset of individuals. Affected individuals were designated mild or classic based on *ATM* mutations and clinical and laboratory features.

**Results:** People with classic A-T were more likely to be younger and IgA deficient and to have higher alpha-fetoprotein levels and lower % forced vital capacity compared to individuals with mild A-T. In classic A-T, the expression of genes required for V(D)J recombination was lower, and the expression of genes required for inflammatory activity was higher. We assigned inflammatory scores to study participants and found that inflammatory scores were highly variable among people with classic A-T and that higher scores were associated with lower *ATM* mRNA levels. Using a cell type deconvolution approach, we inferred that CD4+ T cells and CD8+ T cells were lower in number in people with classic A-T. Finally, we showed that individuals with classic A-T exhibit higher *SERPINE1* (PAI-1) mRNA and plasma protein levels, irrespective of age, and higher *FLT4* (VEGFR3) and *IL6ST* (GP130) plasma protein levels compared with mild A-T and controls.

**Conclusion:** Using an unbiased transcriptomic approach, we identified novel features and developed an inflammatory score to identify subsets of individuals with different inflammatory phenotypes in A-T. Findings from this study could be used to help direct treatment and to track treatment response to therapy.

## Introduction

Ataxia telangiectasia (A-T) is a rare autosomal recessive disease affecting between 1/40,000– 1/300,000 live-births globally (1). Most individuals with A-T reach 3–5 years of age before they begin to exhibit worsening symptoms of the disease, including progressive cerebellar degeneration and ataxia, ocular telangiectasias, and sinopulmonary disease. Individuals with A-T commonly have biallelic heterozygous mutations in the *ATM* gene, which encodes a serine/threonine protein kinase of the same name that is responsible for signaling in response to DNA double-stranded breaks (DSBs) and oxidative stress (2). In people with the classic A-T phenotype, these mutations often lead to a near total loss of functional ATM protein. In the less commonly recognized mild/variant A-T phenotype, individuals often carry *ATM* mutations that result in a moderate reduction in functional ATM protein. These mutations likely account for the milder and later onset A-T phenotype.

Cancer and complications from pulmonary decline are the most common causes of early mortality in people with A-T (3). Current standard of care is supportive and includes monitoring of pulmonary symptoms, weight loss, aspiration, and cancer screening. There are no curative or disease-targeted therapies for A-T that prevent progression of neurodegeneration. Ongoing and completed clinical studies have included monthly infusions of intra-erythrocyte dexamethasone (4) and drug compounds such as metformin (5), nicotinamide adenine dinucleotide (NAD) (6, 7), and N-Acetyl-L-Leucine (8), concluding primarily with negative results. Genetic interventions using antisense oligonucleotides are also being explored in a small set of individuals harboring splice-altering intronic mutations (9).

The lack of mutational hotspots in the *ATM* gene as well as ATM’s role in regulating responses to stochastic DNA DSBs likely contribute to variability of clinical phenotypes. The resulting heterogeneity of clinical trial enrollees may explain inconsistent response to treatment interventions predicted to be effective by preclinical studies. Therefore, identifying peripherally accessible features that can provide mechanistic insights and enhance clinical phenotyping could allow for better assignment of patients to appropriate therapies. In previous studies, we found elevated serum levels of IL-6 and IL-8 in a subset of individuals with A-T (10, 11), consistent with a pro-inflammatory phenotype. In this study we used bulk mRNA-sequencing to broadly characterize the gene expression profiles of PBMCs isolated from individuals with classic and mild A-T. We identified differences in the expression of critical regulatory genes involved in dysregulated biologic processes between classic and mild A-T. We further observed heterogeneous inflammatory activity in A-T-affected individuals and identified novel peripheral protein features based on our bulk RNAseq analysis that may represent biomarker candidates. Findings from this study may be useful for disease classification and for monitoring response to therapy.

## Methods

### Participants

The work described has been carried out in accordance with The Code of Ethics of the World Medical Association (Declaration of Helsinki) for studies involving humans. Participants were recruited from Johns Hopkins University and the Children’s Hospital of Philadelphia. This study was approved by the institutional review board of the Johns Hopkins Medical Institutions (IRB NA_00014314 and NA_00051764) and the institutional review board of the Children’s Hospital of Philadelphia (IRB 20-017524). Participants were recruited from 2018 to 2023. All participants with A-T met the diagnostic criteria for A-T based on clinical symptoms and findings of pathogenic *ATM* mutations and/or elevated alpha-fetoprotein, diminished ATM protein, and increased chromosomal breakage after *in vitro* exposure to x-rays. Demographic information was obtained from chart review. Participants were recruited sequentially from the outpatient Johns Hopkins A-T Clinic and Children’s Hospital of Philadelphia Rare Lung Diseases Frontier Clinic. Participants were assigned to either a mild or classic A-T phenotype based on previously described phenotypic classification strategies (12, 13). Individuals unaffected by A-T were recruited as controls from Johns Hopkins University and the Children’s Hospital of Philadelphia. Control participants answered a brief medical history questionnaire and were excluded from study if they self-identified as being sick at the time of blood draw or as having a severe chronic illness.

### Isolation of peripheral blood mononuclear cells (PBMCs)

PBMCs from peripheral venous blood were isolated with the Ficoll-Paque Plus Method (GE Healthcare Bio-sciences AB) using a process that maintains the viability of B and T lymphocytes to account for relative lymphopenia among A-T versus control participants as previously described (14). Briefly, a 7-mL venous blood sample was transferred into a 50-mL conical tube and PBS was added to achieve a 20-mL volume. 15 mL of Ficoll-Paque Plus was transferred into a separate 50-mL conical tube using a syringe. The 20 mL of diluted blood was gently layered onto the Ficoll. The sample was spun at 400 ×*g* for 30 minutes at room temperature with no brake. The PBMC layer was then collected at the diluted plasma/Ficoll interface. PBS was added to the PBMCs to bring the volume up to 20 mL. The sample was then spun at 200 ×*g* for 10 minutes at room temperature. The supernatant was discarded, and the cells were resuspended in 1 mL of PBS and counted using a TC20 automatic cell counter (Bio-Rad). 4 mL of PBS was added to the sample and spun at 200 ×*g* for ten minutes to remove any contaminating Ficoll, platelets, and plasma proteins. For transcriptional profiling, the sample was resuspended in 350 μL of RLT plus (Qiagen) containing 1% beta-mercaptoethanol, vortexed for 30 seconds, and then transferred to a -80 °C freezer.

### *ATM* genetic variant annotation

Mutations identified in the *ATM* gene during the A-T diagnostic process were reported back to researchers as part of this study. Mutations were annotated with the highest impact calculated molecular consequence using the Ensembl Variant Effect Predictor (VEP) v109.0 (15).

### Bulk RNA sequencing

Total RNA was isolated using the Qiagen AllPrep DNA/RNA Micro kit. Transcriptional profiling was performed as previously described (14, 16) with the following modifications. Libraries for RNA sequencing were prepared with the NEBNext Ultra I RNA Library Prep Kit for Illumina with poly(A) mRNA selection (12-ng input) and sequenced using 1×75 single-end reads on an Illumina NextSeq 2000 instrument using a NextSeq 2000 P2 reagent kit. FASTQ files were generated from read spot intensities in BCL format using BCL2FASTQ v2.17.1.14 with default parameters. Reads were aligned to the NCBI human genome (GRCh38) using the nf-core/rnaseq v3.8.1 Nextflow pipeline (17). In this pipeline, reads were trimmed using TrimGalore!, a wrapper around Cutadapt (18) and FastQC, in order to remove sequencing adapters. Trimmed reads were aligned to the reference genome using STAR 2.7.10a (19) and quantified at the gene level using Salmon 1.9.0 (20) to generate a gene-by-sample counts matrix. Differential gene expression analysis was performed using DESeq2 (21). Differentially expressed genes (DEGs) were identified between groups using the ashr (22) results test with the model design formula *f ∼ cohort + condition*, where *cohort* represents sequencing batch and *condition* is one of unaffected, A-T mild, or A-T classic. DEGs were called with a false-discovery rate (FDR) cutoff of FDR < 0.05 and a minimum log2-fold change (LFC) of magnitude 0.25. Variance stabilized counts used for plotting and tertiary analysis were generated using the rlog function with *blind=FALSE*. Tertiary processing of bulk RNAseq data was performed using scanpy (23). Batch-correction of rlog-normalized counts was performed using Combat-seq (24). Gene set scoring was performed using the hallmark gene sets provided by MSigDB (25) and the JASMINE scoring metric (26). Over-representation analysis (ORA) was performed using gProfiler (27), with runtime options *unordered query*, *only annotated genes*, *g:SCS threshold < 0.05*, and *max term size = 2500*.

### Single-cell RNAseq analysis

Gene count data from (28) were retrieved from the EMBL-EBI ArrayExpress data repository (accession E-MTAB-11452) and processed using scanpy (23). The top 5000 highly variable genes were selected using the *sc.tl.highly_variable_genes* function from scanpy, with *method=”seurat_v3”* and *batch_key=”donor”*. Subsequently, a PCA was performed using the *sc.pp.pca* function with *svd_solver=”arpack”*. This PCA was batch-corrected using the pytorch implementation of the *harmony* algorithm (29) with *batch_key=”donor”*. A cell-cell neighborhood network was generated using the *sc.pp.neighbors* function with *n_neighbors=sqrt(0.25 * adata.n_obs)*, and 2D UMAP was calculated using the *sc.tl.umap* function with *init_pos=”spectral”*. Leiden clustering (30) was performed using the *sc.tl.leiden* function with *resolution=1.25*. Original cell type labels from (28) (*celltype_lvl2_inex_10khvg_reads_res08_new*) were collapsed to parent cell type labels on a per-leiden-cluster basis, and all cells other than B cells, CD4 T cells, CD8 T cells, monocytes, and NK cells were removed.

### Cell type deconvolution of bulk RNAseq data

Cell type proportions for each PBMC bulk RNAseq sample in this study were inferred using CellAnneal (31). Mean counts-per-million (CPM) normalized counts for scRNAseq of each parent cell type were used in order to generate a cell type-specific gene signature matrix. A marker gene dictionary that includes only those genes well-expressed in both the gene signature matrix and bulk RNAseq data was prepared as required using the *cellanneal.make_gene_dictionary* function. Batch-corrected, DEseq2-normalized bulk RNAseq expression data were deconvolved on a per-sample basis with this gene dictionary and gene signature matrix using the *cellanneal.deconvolve* function, with *maxiter=750*.

### Plasma protein concentration measurements

Plasma samples from individuals with A-T and unaffected healthy controls were collected under IRB protocol numbers IRB 20-017524 from Children’s Hospital of Philadelphia and NA_00051764 from Johns Hopkins School of Medicine. We used Abcam *in vitro* SimpleStep ELISA kits (human gp130 (ab246548), Annexin A2 (ab264612), PAI1 (ab269373), and VEGF R3 (ab252350)) to quantify the concentration of plasma proteins of interest. Kits were used according to their individual standard issued protocols. Standard curve dilutions were created from the protein stock provided in the kits and following recommended dilutions. Plasma samples were then diluted 50-fold (PAI1, gp130, VEGF R3) or left undiluted (Annexin A2). Samples were incubated on a pre-coated, 96-well antibody plate with antibody cocktail, shaking for one hour. Wells were washed three times and then incubated for a maximum of 10 minutes with development solution while wrapped in metal foil. Stop solution was immediately added and optical density at 450 nm was read on a Multiskan SkyHigh Microplate Spectrophotometer. A linear standard curve was fitted and used to calculate the protein concentration in each sample.

### Statistical analysis

Statistical testing, beyond that described for bulk RNAseq analysis, was performed using the scipy (32) and statsmodels (33) software packages. Where three or more groups were compared, a Kruskal-Wallis one-way ANOVA on ranks test was performed followed by Dunnett’s tests or one-way Student’s t-tests with Benjamini-Hochberg multiple hypothesis correction as appropriate.

### Data availability

Processed, annotated gene expression data will be provided via the NIH Gene Expression Omnibus (GEO) following peer reviewed publication.

## Results

### Participant phenotype and genotype characteristics

Participants with A-T were recruited from the rare lung disease clinics (CHOP) and multidisciplinary A-T outpatient clinics (Johns Hopkins), both tertiary/quaternary care centers (overview of study design is shown in **Figure 1A**). Individuals with classic A-T accounted for 85% of the A-T participants with a mean age of 11.7 years (**Table 1**). In contrast, the mean age of individuals with mild A-T was 20.6 years, indicative of milder or later onset of disease morbidity in this cohort. In agreement with previously described phenotypic characteristics (12), individuals with classic A-T had higher levels of blood alpha fetoprotein (AFP), higher prevalence of IgA deficiency, and lower forced vital capacity than individuals with mild A-T.

**Figure 1:**
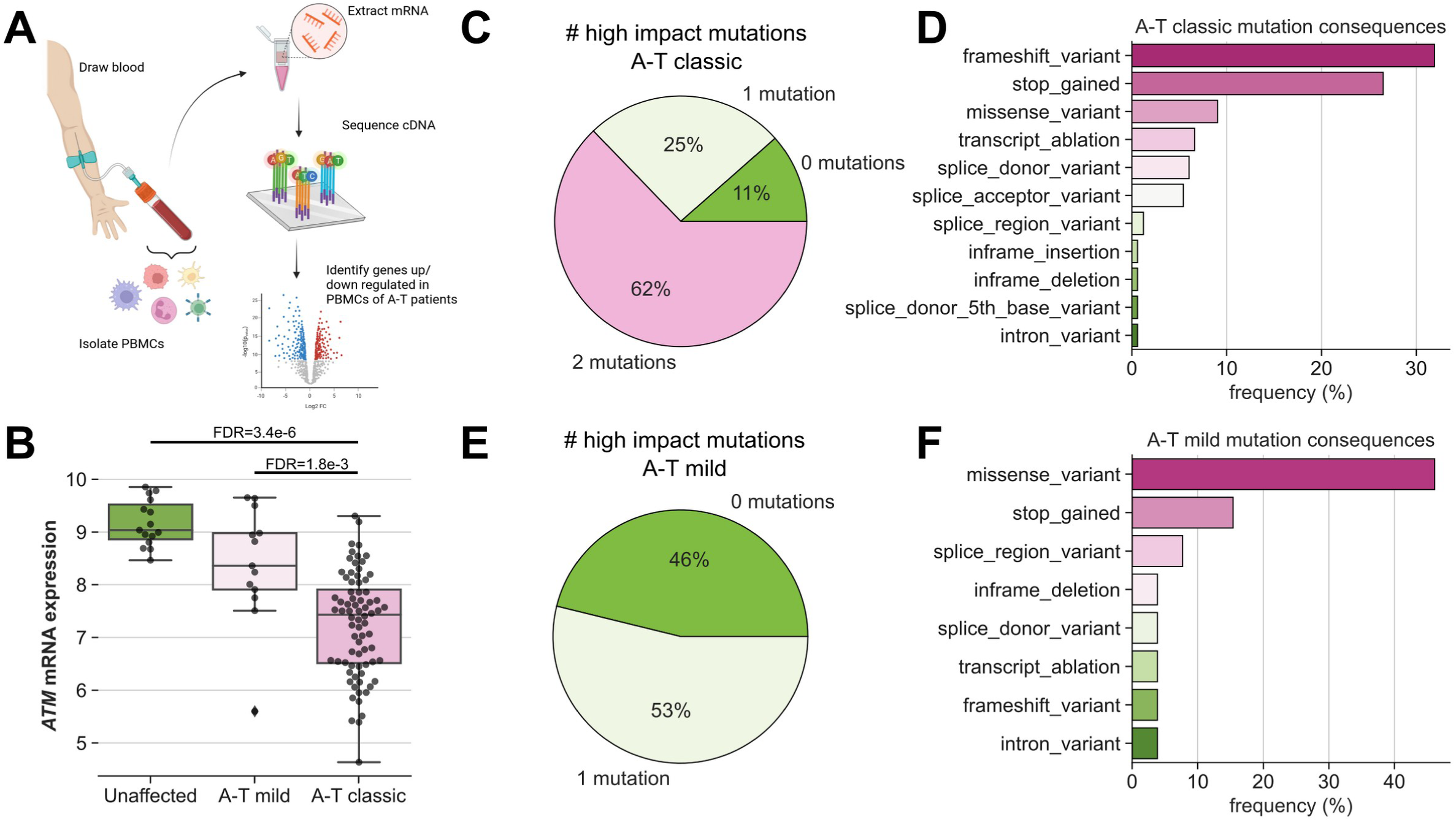
Study design and cohort phenotype/genotype characteristics. **A)** Diagram outlining study design. **B)** Boxplot showing rlog-normalized *ATM* expression in PBMCs stratified by A-T classification, showing decreasing *ATM* mRNA detection with increasing disease severity. **C,E)** Pie charts showing number of mutations with a high impact consequence detected in a given study participant for classic A-T (C) and mild A-T (E) affected individuals. **D,F)** Barplot showing most frequent mutation consequences for mutations detected in classic A-T (D) and mild A-T (F) affected individuals.

**Table 1:** Study participant demographics.

| Mean ± S.D. [Range] | Overall | Classic A-T | Mild A-T | Unaffected controls | P value |
| --- | --- | --- | --- | --- | --- |
| **Study population (n)** | 105 | 77 | 13 | 15 | - |
| **Sex (% female)** | 52.8%<br>(n=55) | 51.3%<br>(n=39) | 53.8%<br>(n=7) | 55.5%<br>(n=9) | 0.98<sup>^</sup> |
| **Age (years)** | 14.6 ±9.1<br>[1-32]<br>(n=105) | 11.7±8.1<br>[1-30]<br>(n=77) | 20.6±8.4<br>[3-32]<br>(n=13) | 24.1±8.1<br>[16-31}<br>(n=15) | **0.0001<sup>^</sup>** |
| **% Forced vital capacity** | 61.9 ±18.8<br>[35-94]<br>(n=47) | 55.9 ±16.4<br>[35-92]<br>(n=37) | 82.8 ±11.0<br>[62-94]<br>(n=10) | - | **0.001<sup>†</sup>** |
| **IgA (% deficient)** | 53%<br>(n=47) | 61%<br>(n=41) | 0%<br>(n=6) | - | **0.004<sup>†</sup>** |
| **AFP (ng/ml)** | 201.3±205.2<br>[2.7-1212.5]<br>(n=82) | 227.1±209.6<br>[21-1212.5]<br>(n=71) | 41.8±45.8<br>[2.7-154.7]<br>(n=11) | - | **0.005<sup>†</sup>** |

<sup>^</sup> Kruskal-Wallis one-way ANOVA on ranks test
<sup>†</sup> One-way Student's t-test

Compared with unaffected controls, individuals with classic or mild A-T each had lower PBMC *ATM* mRNA expression as measured by bulk RNAseq, with individuals with classic A-T exhibiting lower expression compared to individuals with mild A-T (**Figure 1B**). This difference in *ATM* expression was associated with *ATM* genotype. Individuals with classic A-T were more likely to have bi-allelic high-impact *ATM* mutations (**Figure 1C**), including frameshift, stop-gain, and transcript ablation variants (**Figure 1D**) that would severely impact the ability to detect *ATM* mRNA due to increased nonsense mediated decay. Individuals with mild A-T were more likely to have mono-allelic or no high impact *ATM* mutations (**Figure 1E**), presenting predominantly with missense variants (**Figure 1F**). No individuals with mild A-T had bi-allelic high-impact *ATM* mutations.

### PBMC bulk RNAseq reveals dysregulation of genes involved in V(D)J recombination and inflammation in individuals with classic A-T

Bulk RNAseq was performed on RNA isolated from PBMCs of individuals with classic A-T, mild A-T, and unaffected controls. Differential gene expression (DEG) testing indicated that 1011 genes were significantly higher in expression and 1060 genes were significantly lower in expression in individuals with classic A-T compared with unaffected controls (**Figure 2A, Supplemental Table 1**). Only 12 genes were significantly higher in mild A-T individuals relative to unaffected controls, indicative of similarities in the gene expression profiles of PBMCs from these two groups, while the expression 86 genes was higher and 335 genes was lower when comparing individuals with classic versus mild A-T (**Supplemental Figure 1A-C, Supplemental Table 2**).

**Figure 2:**
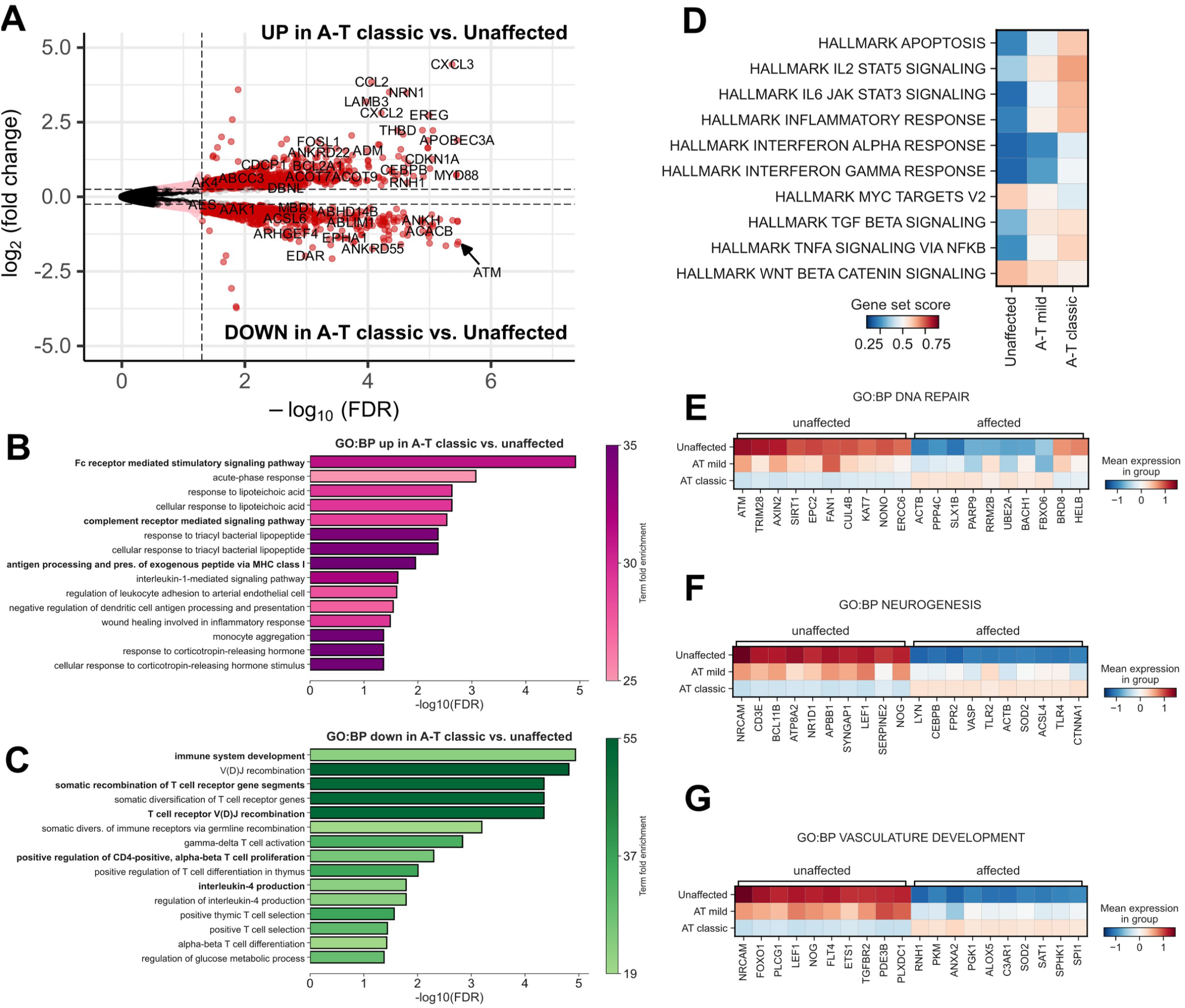
Bulk RNAseq of PBMCs in classic A-T suggests loss of T-cell V(D)J recombination activity and enrichment in monocyte-associated inflammatory pathways. **A)** Volcano plot showing differences in gene expression in PBMCs from classic A-T vs. unaffected control individuals. **B,C)** Over representation analysis (ORA) of GO Biological Processes (GO:BP) for genes increased (B) and decreased (C) in expression in classic A-T vs. unaffected control individuals. **D)** Matrixplot of selected MSigDB hallmark geneset scores. **E,F,G)** Matrix plots showing mean Z-scored expression of the differentially expressed genes involved in the GO:BPs DNA repair (E), neurogenesis (F), or vascular development (G).

We performed an over-representation analysis (ORA) on DEGs by comparing individuals with classic A-T to unaffected controls. Using the GO:Biological Process ontology (GO:BP), we found a significant enrichment in pathways involved in a variety of monocyte-driven inflammatory processes, including *Fc receptor mediated signaling*, *complement receptor mediated signaling*, and *antigen processing/presentation via MHC class I* in individuals with classic A-T compared to unaffected controls (**Figure 2B**). Conversely, pathways associated with T-cell specific processes, including *somatic recombination of T cell receptor gene segments*, *T cell receptor V(D)J recombination*, and *positive regulation of CD4 T cell proliferation* were significantly enriched in unaffected controls versus individuals with classic A-T (**Figure 2C**). These ORA results correlated well with the results of MSigDB hallmark gene set scoring, which indicated that pathways involved in inflammatory responses are largely enriched in both mild A-T and classic A-T (**Figure 2D**). These data indicate that PBMCs from A-T-affected individuals have lower expression of genes required for T-cell development and function and higher expression of inflammatory monocyte-related genes, providing evidence for a dysregulation of systemic inflammatory activity in individuals with A-T.

### Genes involved in DNA damage repair, neurogenesis, and vascular development are dysregulated in PBMCs from A-T-affected individuals

We further asked which genes involved in critical biological processes affected by A-T were differentially expressed in PBMCs from A-T-affected individuals relative to unaffected controls; namely those genes related to DNA repair (**Figure 2E**), neurogenesis (**Figure 2F**), and vascular development (**Figure 2G**). Across genes involved in the DNA repair process, we found that expression of *AXIN2*, a gene encoding an inhibitor of the WNT signaling pathway (34), was lower in classic A-T relative to unaffected controls, as was *ERCC6*, which encodes a critical mediator of ATM, CHEK2-dependent DNA repair processes (35). We likewise found that *UBE2A* and *FBXO6* were both higher in expression in individuals with classic A-T, which may represent a compensatory response to ATM loss, as *UBE2A* encodes a protein required for replicative DNA repair (36), and *FBXO6* encodes a protein that identifies and promotes the ubiquitination of activated CHEK1 (37), which is phosphorylated by functional ATM in response to DNA damage (38). Taken together, these data indicate that several genes involved in DNA damage response, neurogenesis, and vascular development are dysregulated in the PBMCs of individuals with classic A-T and therefore may be involved in the development of the hallmark cerebellar atrophy and telangiectasia development found in A-T-affected individuals.

### Heterogeneous expression of inflammatory response genes observed in individuals with classic A-T

Observing greater expression of genes involved in typical inflammatory response-related processes during cohort-level DEG analysis, we wondered whether the variance across individuals with either classic or mild A-T in this regard was narrow or broad. We plotted a kernel density estimate of the calculated hallmark inflammatory response score (**Figure 2B**) split across each group and noted heterogeneity within each group despite different between-group mean scores (**Figure 3A**). In both the classic and mild A-T groups, two peaks in these distributions were observed, and we used the approximate location of these peaks to set thresholds between groups of participants with low, moderate, and high inflammatory response categories. The proportion of participants within each inflammatory response level varied by group when compared across the unaffected control, mild, and classic A-T groups (**Figure 3B**), with similar proportions of participants in the high inflammatory response group in both the mild and classic A-T groups but with an expansion in the size of the moderate inflammatory response category in the classic A-T group.

**Figure 3:**
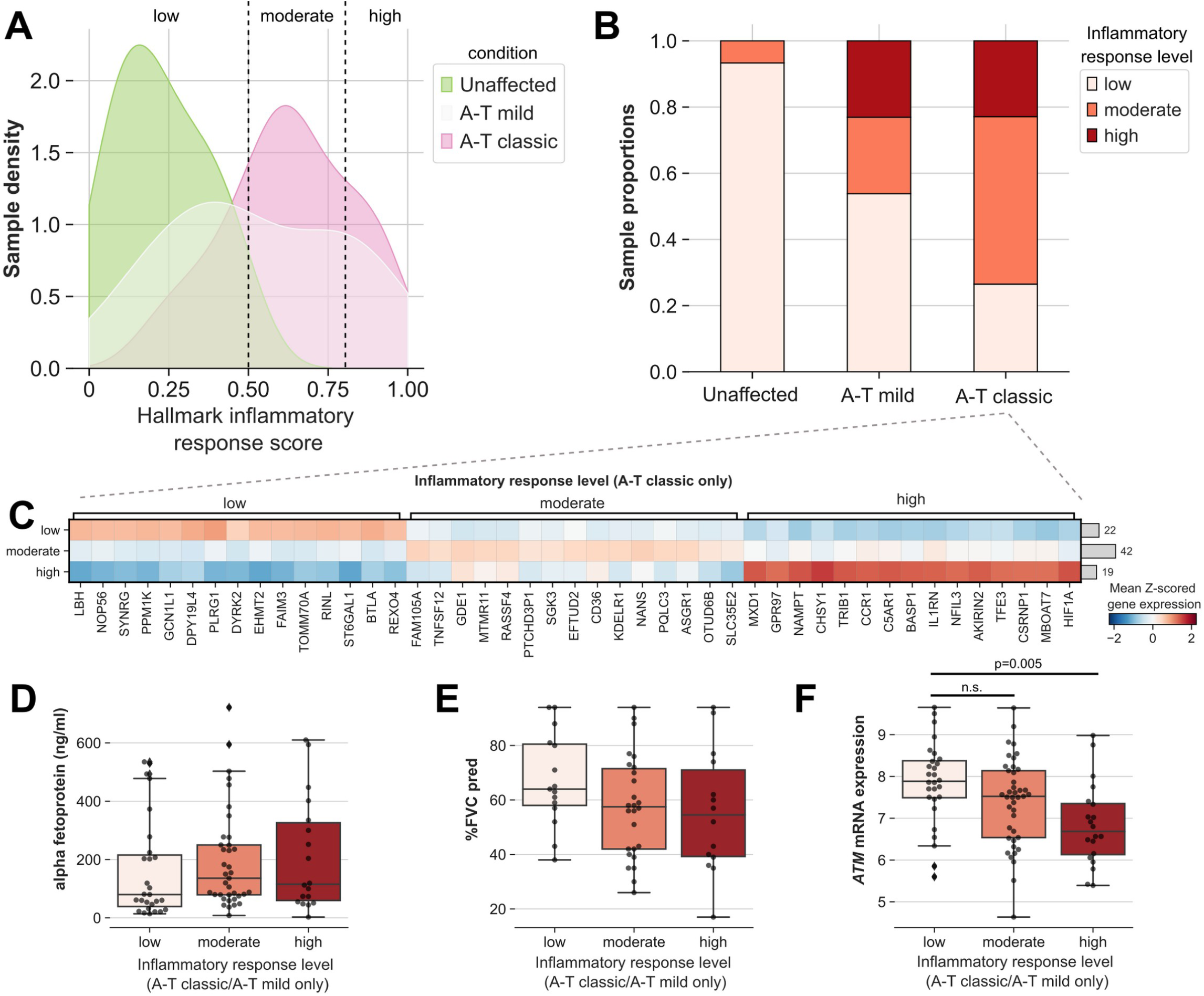
A-T classic affected individuals show heterogeneous inflammatory pathway activities. **A)** Kernel density estimate (KDE) plot showing distribution of hallmark inflammatory response scores stratified by condition. **B)** Stacked barplot showing proportion of samples with low [0,0.5), moderate [0.5, 0.8), or high [0.8, 1] inflammatory response levels, based on scores in (A). **C)** Matrixplot showing expression of genes differentially expressed across inflammatory response levels in classic A-T affected individuals only. **D,E,F)** Boxplots of measured alpha-fetoprotein (D), % forced vital capacity (E), and rlog-normalized *ATM* mRNA expression (F) in mild A-T and classic A-T affected individuals, stratified by inflammatory response level.

This heterogeneity across low, moderate, and high inflammatory response levels among individuals with classic A-T prompted us to ask whether changes in gene expression in the PBMCs of these individuals were identifiable across inflammatory response levels. Many genes were higher in expression in individuals with classic A-T and a high inflammatory response level are involved in inflammatory pathways or are predominantly expressed in monocytes (**Figure 3C**). Of the genes that were lower in expression in individuals with classic A-T and a high inflammatory response level, we identified *LBH* as an important feature, as loss of *LBH* induces S-phase arrest and failure to progress through the cell cycle, in turn leading to worsened inflammation in certain contexts such as mouse models of rheumatoid arthritis (39). *ATM* mRNA expression within the mild and classic A-T groups decreased with increasing inflammatory response level (**Figure 3F**); however, no significant changes were observed in blood alpha fetoprotein levels (**Figure 3D**) or percent forced vital capacity (**Figure 3E**). Taken together, these data indicate that there is evidence of significant heterogeneity in the inflammatory response in individuals affected with classic A-T.

### Loss of CD4+ and CD8+ T cells in A-T-affected individuals inferred using bulk RNAseq cell type deconvolution

Changes in the expression of PBMC subtype-specific genes as measured by bulk RNAseq may arise due to gene regulatory mechanisms that modify gene expression activity across all/many cells or by changes in the proportion of specific cell subtypes expressing that gene that were present in the sample at the time of mRNA extraction. Using a reference single-cell RNAseq atlas of human PBMCs (28), we aimed to infer the proportion of different major PBMC subtypes in each of our bulk RNAseq samples in order to investigate whether changes in cell type proportions were a potential explanation for our results.

After re-processing the scRNAseq data and subsetting to include subtypes of B cells, CD4 T cells, CD8 T cells, monocytes, and NK cells (**Figure 4A**), we used CellAnneal (31) to identify robustly expressed marker genes for each parent cell type (**Figure 4B**) in order to infer cell the proportions of each parent cell type within each bulk RNAseq sample. Compared to samples from unaffected controls, samples from individuals with mild and classic A-T exhibited a lower relative proportion of CD4+ and CD8+ T cells, with a corresponding expansion of the inferred monocyte fraction (**Figure 4D**), corresponding with previous reports (40) (41) (42). Within samples from individuals with classic A-T and splitting across inflammatory response levels, we observed lower CD4+ and CD8+ T cell proportions in samples with moderate and high inflammatory response scores compared to samples with low inflammatory response scores, showing marked variability within the classic A-T group (**Figure 4E**).

**Figure 4:**
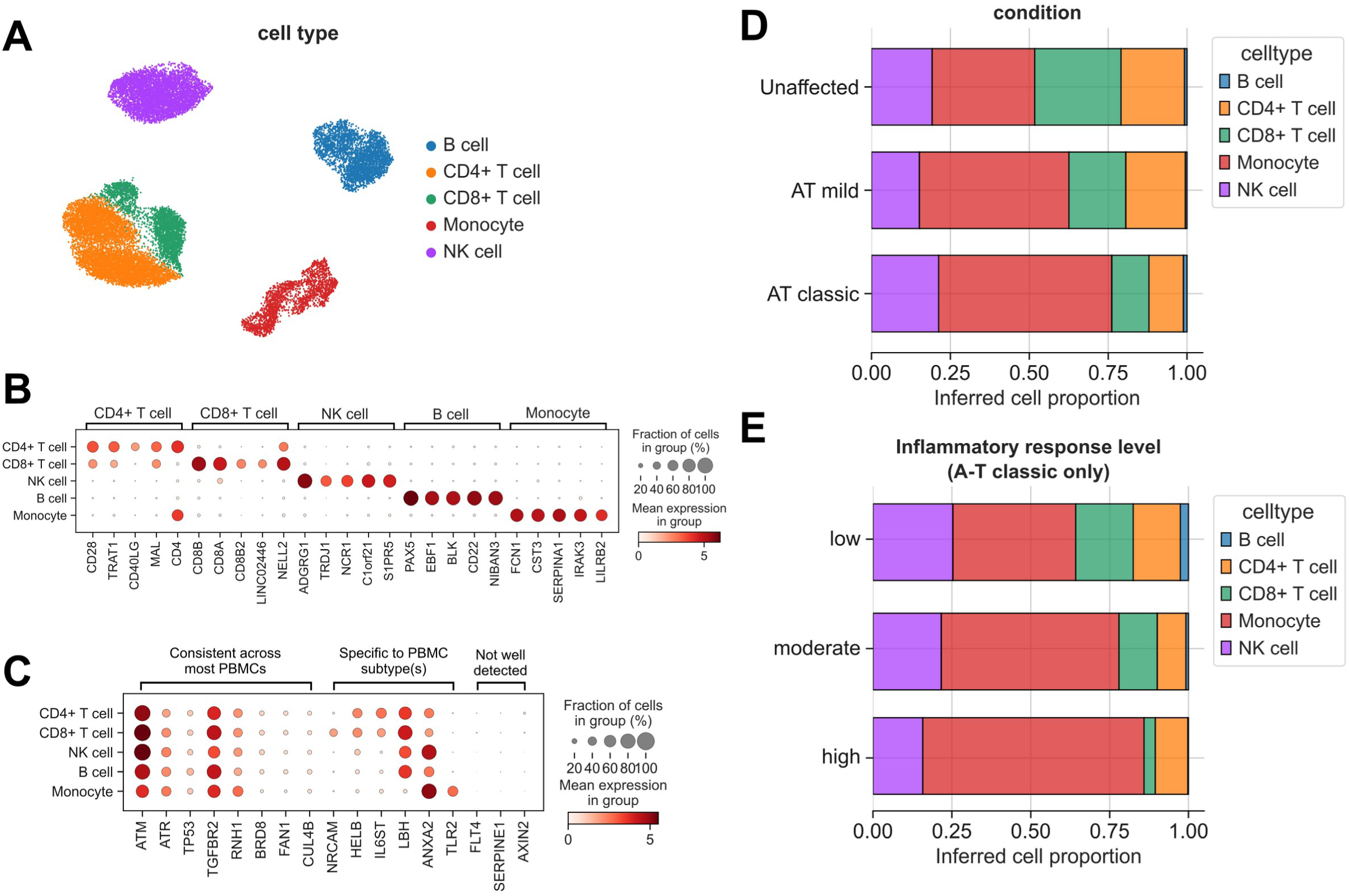
Inference of cell type proportions from bulk RNAseq indicates loss of CD4+/CD8+ T-cell populations in A-T affected individuals. **A)** Uniform Manifold Approximation and Projection (UMAP) plot of subset of data from (Hagemann-Jensen et. al., 2022) Smart-seq3xpress atlas of human PBMCs, showing parent cell type labels mapped here for deconvolution. **B)** Dotplot showing expression of a subset of identified marker genes across parent cell type groups. **C)** Dotplot showing expression of A-T related genes of interest in this study across parent cell type groups. **D)** Stacked barplot of CellAnneal-inferred cell type proportions in bulk RNAseq samples stratified by condition. **E)** As in (D) but for classic A-T affected individuals only, stratified by inflammatory response level.

As large changes in PBMC subtype proportions may be a contributor to changes in gene expression measured by bulk RNAseq, we used this reference scRNAseq atlas to measure the expression of certain genes of interest across PBMC subtypes (**Figure 4C**). Multiple disease-related genes, such as *ATM, ATR,* and *TP53* were expressed at similar levels across all PBMCs. Others, such as *NRCAM, LBH,* and *SERPINE1*, were highly variable across PBMC subtypes or not well detected in this scRNAseq dataset. Taken together, these results indicate that CD4+ T cell/CD8+ T cell loss is a partial contributor to gene expression differences observed in these bulk RNAseq data, highlighting the heterogeneity of immunologic features across individuals affected with classic A-T and providing a framework for identifying gene dysregulation in both PBMC subtype-specific and non-specific contexts.

### Identification of PAI-1 (SERPINE1) as a plasma protein correlated with A-T severity

Having observed differences in the transcriptomes of PBMCs from unaffected and A-T-affected individuals, we were interested to determine whether any differentially expressed genes might be detectable at the protein level in blood plasma. Hence, we used the Human Protein Atlas’ blood secretome dataset (43) to filter our bulk RNAseq data to include only those genes detectable by either protein ELISA or protein mass-spectrometry in blood plasma and then re-performed differential gene expression testing (**Figure 5A**). Comparing unaffected controls to individuals with classic A-T, we detected significantly lower expression of genes such as *IL6ST, FLT4*, and *CFH* as well as greater expression of *TIMP1, IL1RN*, and *SERPINE1*.

**Figure 5:**
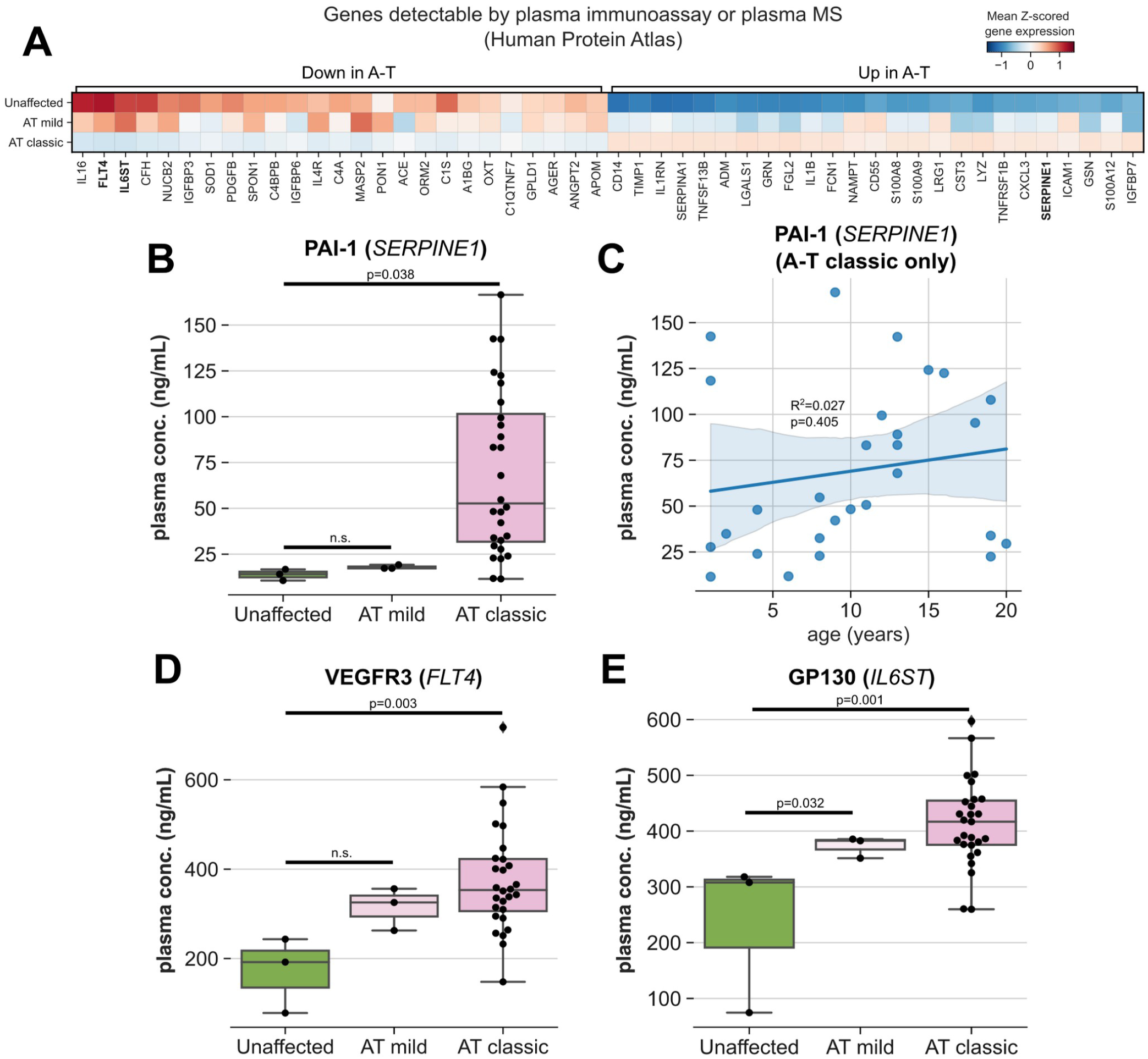
Identification and characterization of protein biomarkers of A-T disease severity. **A)** Matrixplot of expression of differentially expressed genes across classic A-T and unaffected individuals, restricted to those genes whose protein products are detectable by blood plasma immunoassay or mass-spectrometry according to the Human Protein Atlas. **B,D,E)** Boxplots of protein concentrations in blood plasma by ELISA for PAI-1 (*SERPINE1*) (B), VEGFR3 (*FLT4*) (D), and GP130 (*IL6ST*) (E), stratified by condition. **C)** Linear regression of PAI-1 plasma concentration versus age in individuals with classic A-T.

We measured blood plasma protein concentrations of VEGFR3 (*FLT4*) due to its role in regulating angiogenesis (44), GP130 (*IL6ST*) due to its role in modulating immune responses (45), and PAI-1 (*SERPINE1*) due to its association with premature aging phenotypes (46, 47) and its putative biochemical links to ATM DNA repair function (48, 49). PAI-1 concentrations were significantly higher in plasma from individuals with classic A-T relative to both unaffected controls and individuals with mild A-T (**Figure 5B**). PAI-1 protein concentrations did not correlate with age (**Figure 5C**), indicating that PAI-1 concentration may be more suitable for tracking disease progression/response to treatment over time than AFP concentration, which has been shown to increase with age (50). Despite lower expression of *FLT4* and *IL6ST* mRNA in PBMCs from individuals with classic A-T, VEGFR3 and GP130 blood plasma concentrations were higher in plasma from individuals with classic A-T relative to unaffected controls (**Figure 5D**). These findings suggest that these plasma proteins are derived mostly from a non-PBMC source.

## Discussion

A-T presents a wide range of challenges for individuals affected by the disease due to its multi-system presentation. While preliminary work on the development of genetic therapies targeting a small subset of mutations in the *ATM* gene has recently been described (9), the development of therapies that may have broad utility has been hindered by the lack of molecular insights that relate to development and progression of classic A-T disease manifestations. In addition, the lack of robust, accessible peripheral biomarkers that might be used to characterize disease severity and monitor response to therapy, especially considering the heterogeneity of phenotypic presentation in A-T, has perhaps been underappreciated.

To address these challenges in this study, we performed bulk RNAseq on PBMCs isolated from individuals affected with A-T and from unaffected controls. We demonstrated that PBMCs from A-T-affected individuals exhibit significantly lower expression of genes involved in V(D)J recombination and T cell development, instead showing greater expression of monocyte-specific inflammatory genes. Generating an inflammatory response score using hallmark genesets from MSigDB and reanalyzing a previously published scRNAseq atlas of PBMCs, we performed cell type deconvolution and inferred a loss of CD4+ T-cells and CD8+ T-cells in A-T affected individuals relative to unaffected controls. We further demonstrated that inflammatory response scores and inferred T cell losses are heterogeneous in individuals with classic A-T. Finally, we leveraged data from the Human Protein Atlas to identify differentially expressed genes in our bulk RNAseq data that are detectable in human blood plasma and showed that PAI-1 (*SERPINE1*) protein concentration is significantly higher in blood plasma from individuals with classic A-T relative to both individuals with mild A-T and unaffected controls and, importantly, does not correlate with age.

Genes involved in multiple A-T disease-related processes, including DNA repair, neurodevelopment, and vasculogenesis were dysregulated in the PBMCs of A-T affected individuals. As we have previously shown (14), *NRCAM* mRNA was significantly lower in expression(14, 51). As classic A-T is characterized by cerebellar degeneration and abnormal saccadic eye movements, this finding suggests there may be an inverse association between expression of NRCAM and these A-T phenotypes. This potential association is further evidenced by data indicating that *NRCAM* is regulated by p53, a phosphorylation target of ATM that serves as a master regulator of cell cycle progression, DNA repair, and apoptosis (52). However, *NRCAM* is also detected in CD8+ T cells in at least one reference PBMC scRNAseq atlas (28) (**Figure 4C**), indicating that observed differences in *NRCAM* expression may be due to overall CD8+ T cell loss in our study. Interestingly, lower expression of *NRCAM* has also been reported in lymphoblastoid lines from patients with temporal lobe dementia due to mutations in *MAPT* (*53*). We found that other genes, such as *PLXDC1*, which is involved in endothelial capillary morphogenesis (54), and *TGFBR2*, which encodes an important receptor for the TGF-β family of ligands (55), were lower in expression in individuals with classic A-T. TGF-β signaling is a critical regulator of cell proliferation and differentiation in many contexts (56), and genetic variants in genes related to this pathway have been linked to the development of hereditary hemorrhagic telangiectasias (57) and developmental neurovascular defects (58). Interestingly, ATM is responsible for stabilizing TGFBR2 in response to DNA damage (59), indicating that the decrease in *TGFBR2* expression in the PBMCs from A-T affected individuals may be directly related to loss of ATM function, especially given that *TGFBR2* is expressed in all PBMC subtypes (**Figure 4C**). Some genes involved in these biological processes were higher in expression in A-T-affected individuals, including *RNH1*, an RNase inhibitor which also inhibits angiogenin (60), and *TLR2/TLR4*, genes encoding toll-like receptors essential for innate immune response (61) and additionally involved in neural differentiation (62) and modulation of neural responses (63).

We identified multiple plasma proteins, including VEGFR3 (*FLT4*), GP130 (*IL6ST*), and PAI-1 (*SERPINE1*) that represent putative features that correlate with A-T severity. While loss of *FLT4* expression was observed in the PBMCs of individuals affected by A-T, we detected higher concentrations in plasma of its protein product, VEGFR3. These results suggest that plasma VEGFR3 protein derives from an alternative source, likely endothelial cells that use VEGFR3 to modulate vascular permeability and sprouting angiogenesis (44, 64).

By both bulk RNAseq and protein ELISA, expression of *SERPINE1* (PAI-1) was higher in individuals with classic A-T compared to both unaffected controls and individuals with mild A-T. Lower levels of PAI-1 have been associated with protection against biological aging (46), and increased expression has likewise been linked to diseases with premature aging phenotypes such as Hutchinson-Gilford progeria syndrome (47). The association between *SERPINE1*/PAI-1 expression and loss-of-function *ATM* mutations is worthy of further study, as the product of *SERPINE2*, a paralog of *SERPINE1*, directly interacts with ATM and modulates its function (48). Both PAI-1 and the product of *SERPINE2* share the same ATM binding domain, so greater PAI-1 expression in individuals with classic A-T may indicate a compensatory response to loss of functional ATM. Treatment of human adipocytes with troglitazone has been shown to decrease PAI-1 expression (65), suggesting that PAI-1 may be a druggable target in A-T if functional links to ATM and related A-T pathology can be established.

In this study, we examined gene expression from PBMCs in individuals with classic and mild A-T. Using an unbiased transcriptomic approach, we identified novel features and developed an inflammatory score to identify subsets of individuals with different inflammatory phenotypes. We also identified several plasma proteins differentially regulated in patients with classic A-T. Findings from this study could be useful in directing treatment and tracking treatment response to therapy.

