## Supplemental figures and legends for "Transcriptional profiling of peripheral blood mononuclear cells identifies inflammatory phenotypes in ataxia telangiectasia"

**Supplemental Figure 1:** Bulk RNAseq of PBMCs reveals defects in T cell differentiation marker genes in individuals with classic relative to mild A-T. A) Volcano plot showing changes in gene expression in PBMCs from classic A-T versus mild A-T affected individuals. B,C) Over representation analysis (ORAs) of GO Biological Processes (GO:BP) for genes increased (B) and decreased (C) in expression in classic A-T vs. mild A-T affected individuals.

**Supplemental Table 1:** Differential gene expression testing results comparing classic A-T to unaffected controls. Related to Figure 2A-C.

**Supplemental Table 2:** Differential gene expression testing results comparing classic A-T to mild A-T. Related to Supplemental Figure 1.

**Supplemental Figure 1**

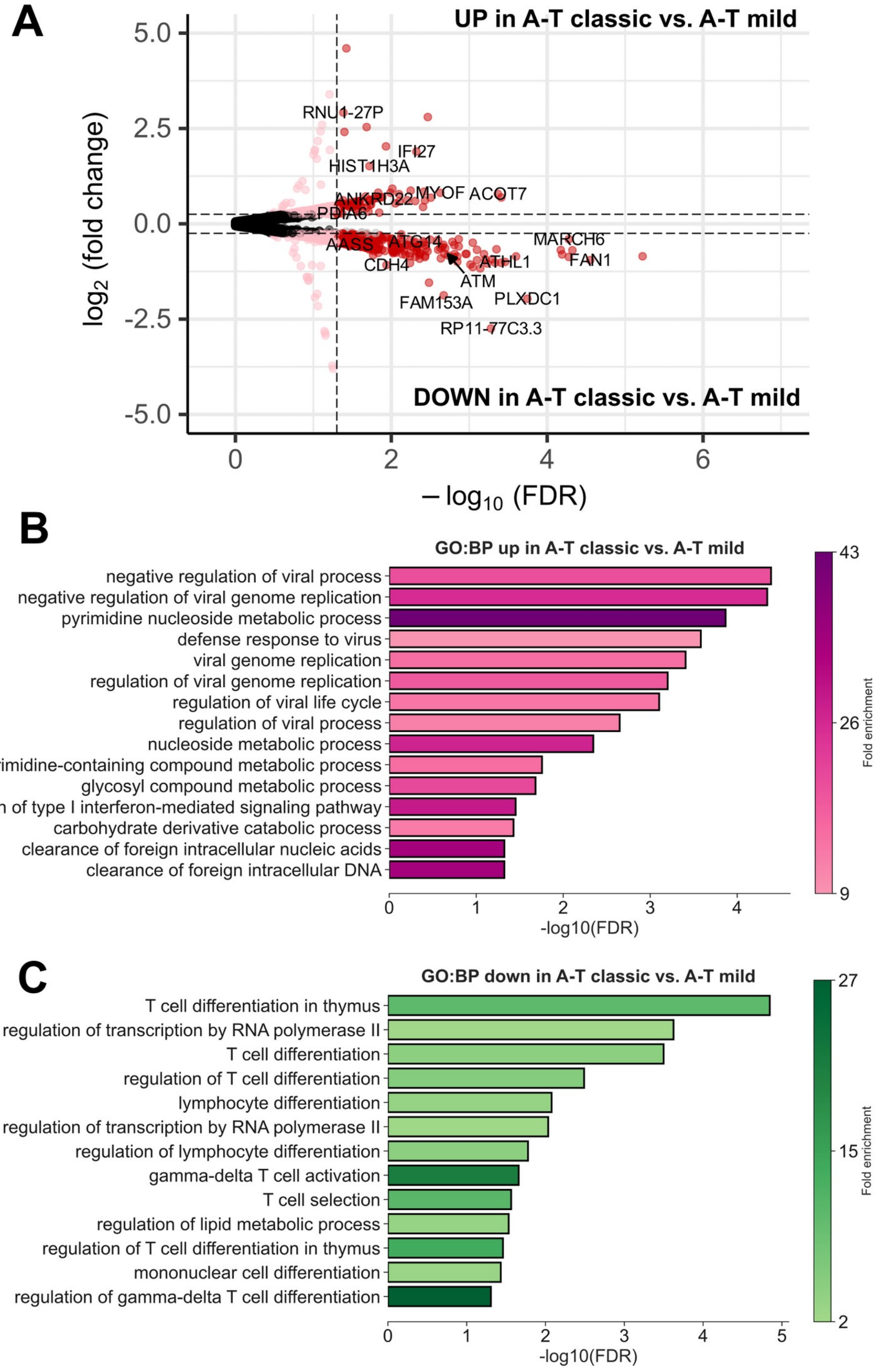
